# Oxidized thioredoxin-1 restrains the NLRP1 inflammasome

**DOI:** 10.1101/2021.09.20.461118

**Authors:** Daniel P. Ball, Alvin E. Wang, Charles D. Warren, Qinghui Wang, Andrew R. Griswold, Sahana D. Rao, Daniel A. Bachovchin

## Abstract

At least six human proteins detect danger-associated signals, assemble into complexes called inflammasomes, and trigger pyroptotic cell death. NLRP1 was the first protein discovered to form an inflammasome, but the danger signals and molecular mechanisms that control its activation have not yet been fully established. Here, we report that the NACHT-LRR region of NLRP1 directly binds to oxidized form of thioredoxin-1 (TRX1). We found that NLRP1 requires the ATPase activity of its NACHT domain to associate with TRX1, and that this interaction represses inflammasome activation. Moreover, we discovered that several patient-derived missense mutations in the NACHT-LRR region of NLRP1 weaken TRX1 binding, leading to inflammasome hyperactivation and autoinflammatory disease. Overall, our results establish that oxidized TRX1 binds to and restrains the NLRP1 inflammasome, thereby revealing a link between the cellular redox environment and innate immunity.

## Main Text

The human NLRP1 (nucleotide-binding domain leucine-rich repeat pyrin domain-containing 1) protein has an N-terminal pyrin domain (PYD), followed by a disordered region and nucleotide-binding (NACHT), leucine-rich repeat (LRR), function-to-find (FIIND), and caspase activation and recruitment (CARD) domains (Fig. 1A). CARD8 is the only other human protein with a FIIND, but it lacks the N-terminal structured domains (Fig. 1A). Both NLRP1 and CARD8 undergo autoproteolysis within their FIINDs, generating autoinhibitory N-terminal (NT) and inflammatory C-terminal (CT) fragments that remain associated (*1-3*). The proteasome-mediated degradation of the NT fragments releases the CT fragments from autoinhibition (*4, 5)*. Initially, each freed CT fragment is restrained in a ternary complex with one copy of the full-length (FL) sensor protein (NLRP1 or CARD8) and one copy of either dipeptidyl peptidase 8 or 9 (DPP8/9) (*6, 7*). However, if enough CT fragments are released or if the ternary complex is disrupted by DPP8/9-binding ligands, the CT fragments assemble into caspase-1-activating structures called inflammasomes and trigger pyroptosis.

**Fig. 1.**
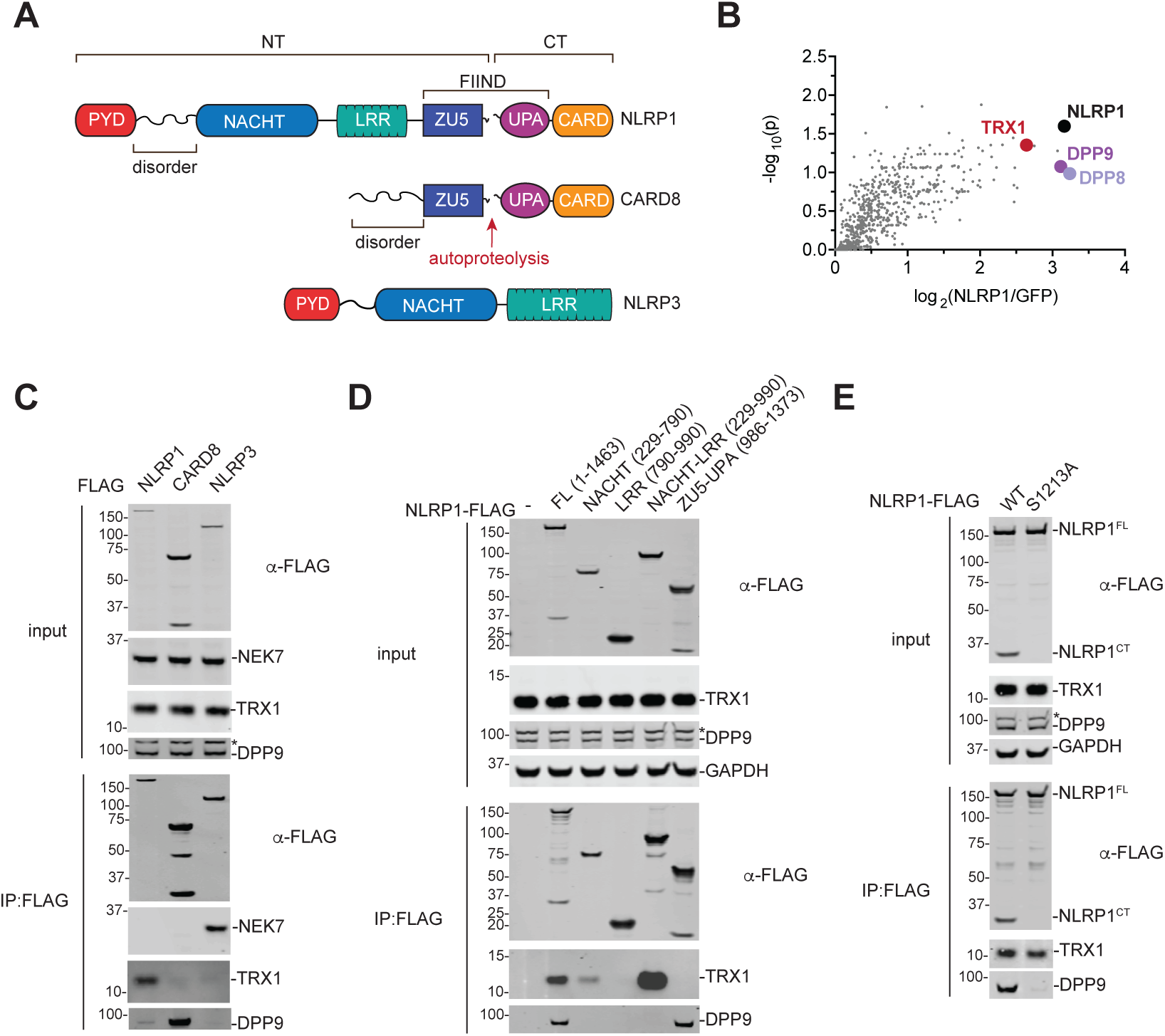
The NACHT-LRR region of NLRP1 associates with TRX1. (**A**) Schematic of the human NLRP1, CARD8, and NLRP3 proteins. The FIINDs of NLRP1 and CARD8 undergo autoproteolysis between the ZU5 and UPA subdomains. NT, N-terminus. CT, C-terminus. (**B**) HEK 293T cells were transfected with GFP- or NLRP1-FLAG. Lysates were subjected to anti-FLAG IP and quantitative MS analyses. The volcano plot depicts proteins enriched in the NLRP1-FLAG IP. (**C**-**E**) HEK 293T cells were transiently transfected with the indicated FLAG-tagged constructs, subjected to anti-FLAG IP, and analyzed by immunoblotting. In **D**, numbers indicate amino acid residues. An asterisk (*) denotes a background band.

At least three seemingly unrelated agents – the enteroviral 3C protease (*8, 9*), long double-stranded RNA (dsRNA) (*10*), and DPP8/9 inhibitors (*11-16*) – activate the human NLRP1 inflammasome. 3C protease cleaves within the disordered linker between the PYD and NACHT domains, creating an unstable neo-N-terminus that is degraded by the N-end rule proteasome pathway. dsRNA directly interacts with the NACHT-LRR region, triggering ATP hydrolysis, and presumably NT degradation and CT release, via unknown mechanisms. DPP8/9 inhibitors, including Val-boroPro (VbP), both accelerate the proteasome-mediated degradation of the NT fragment through a poorly characterized pathway and destabilize the NLRP1^FL^-NLRP1^CT^-DPP8/9 repressive ternary complex (*6, 7, 14, 16, 17*). Of these three triggers, only DPP8/9 inhibitors also activate the rat and mouse NLRP1 inflammasomes (*15*) and the human CARD8 inflammasome (rodents do not express CARD8) (*13, 18*). As such, only DPP8/9 inhibitors, and not dsRNA and 3C protease, are likely related to the primordial function of the NLRP1 inflammasome (*19*).

The NLRP3 inflammasome sensor protein is structurally similar to the NLRP1 NT fragment (Fig. 1A). Notably, the mitotic kinase NEK7 directly interacts with NLRP3’s NACHT-LRR region to license inflammasome activation (*20-22*). We reasoned that an endogenous protein might similarly bind to N-terminal domains of NLRP1. We therefore performed an immunoprecipitation-mass spectrometry (IP-MS) experiment in HEK 293T cells with FLAG-tagged NLRP1 or FLAG-tagged GFP to identify potential NLRP1 binding partners (Fig. 1B, Table S1). This experiment not only identified DPP8 and DPP9 as NLRP1-binding proteins, as expected, but also revealed that thioredoxin-1 (TRX1) potentially interacted with NLRP1 as well (Fig. 1B). We confirmed that TRX1 indeed co-immunoprecipitated human NLRP1, but not CARD8 or NLRP3, by immunoblotting (Fig. 1C). Moreover, we found that TRX1 also interacted with all functional rat and mouse NLRP1 alleles (Fig. S1A,B), showing that TRX1 binding, like DPP8/9 binding, is an evolutionarily conserved function of NLRP1.

We next expressed various truncated forms of FLAG-tagged NLRP1 to identify the region that interacts with TRX1. We found that the isolated NACHT domain retained some affinity for TRX1, but the isolated LRR and FIIND domains did not (Fig. 1D). However, the entire NACHT-LRR region bound more strongly to TRX1 than the isolated NACHT, indicating that these two domains together mediate this protein-protein interaction. DPP9 associates with the NLRP1 FIIND (*6, 7, 14*), and, as expected, only the full-length NLRP1 protein and isolated FIIND bound to DPP9. Thus, TRX1 and DPP9 bind to distinct regions of NLRP1. Consistent with these distinct binding regions, autoproteolysis-deficient NLRP1 (S1213A), which has dramatically impaired binding to DPP9, retained binding to TRX1 (Fig. 1E).

TRX1 is a ubiquitous, highly conserved oxidoreductase that maintains redox balance in the cytosol and protects proteins from oxidative damage and aggregation. TRX1 uses a cysteine (Cys32) within a conserved CGPC motif (residues 32-35) to react with disulfide bonds in oxidized proteins. The second cysteine (Cys35) then resolves the disulfide bond between the target protein and TRX1, thereby reducing the target protein and oxidizing TRX1. The disulfide bond in oxidized TRX1 is then reduced by thioredoxin reductase-1 (TXNRD1), which uses NADPH as an electron donor. To determine if Cys32 and/or Cys35 were critical for binding to NLRP1, we expressed FLAG-tagged WT, C32S, C35S, or C32S/C35S TRX1 together with untagged NLRP1 before performing an anti-FLAG IP. We found that both Cys32 and Cys35 were required for binding (Fig. 2A). It should be noted that NLRP1 and TRX1 could, in theory, form a stable intermolecular disulfide bond. However, this does not occur, as the C35S mutation, which would trap such a bond, dramatically weakened the NLRP1-TRX1 interaction.

**Fig. 2.**
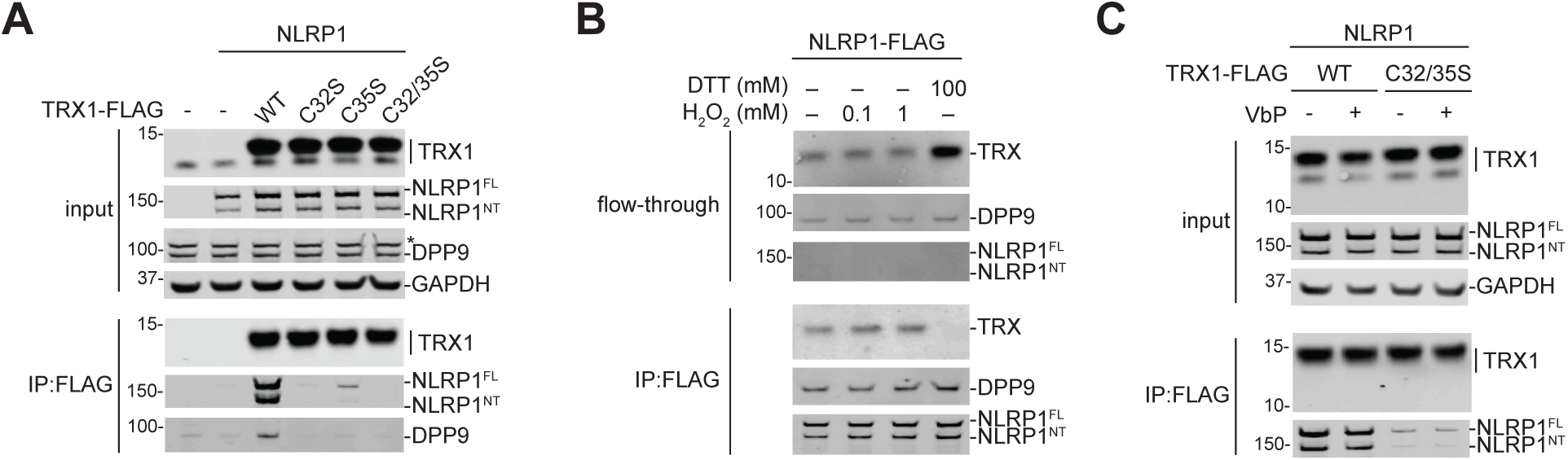
Only oxidized TRX1 binds NLRP1. (**A-C**) HEK 293T cells were transfected with the indicated constructs before lysates were harvested and subjected to anti-FLAG IP and immunoblotting analysis. Anti-FLAG beads were treated with the indicated concentrations of (anti)oxidants in **B** and VbP (10 µM) in **C**. The flow-through in **B** is the unbound fraction after (anti)oxidant treatment. An asterisk (*) denotes a background band.

Instead, we hypothesized that NLRP1 was exclusively binding to either the reduced or the oxidized form of TRX1. To determine which state of TRX1 was bound to NLRP1, we immobilized FLAG-tagged NLRP1 protein from HEK 293T lysates on anti-FLAG beads, and then treated these beads with the oxidizing agent hydrogen peroxide (H2O2) or the reducing agent dithiothreitol (DTT) (Fig. 2B, Fig. S2A). We found that H2O2 slightly strengthened the TRX1 interaction, and conversely that DTT abolished the interaction. Similarly, we found that reduced glutathione (GSH) dramatically weakened the interaction as well (Fig. S2A). Thus, only oxidized TRX1 (i.e., the disulfide form of TRX1) binds to NLRP1. Notably, neither H2O2 nor DTT affected the NLRP1-DPP9 interaction (Fig. 2B). Moreover, the DPP8/9 inhibitor VbP weakened the NLRP1-DPP9 interaction in this assay (Fig. S2B), as expected, but had no direct impact on the NLRP1-TRX1 interaction (Fig. 2C, Fig. S2C). The dsRNA analog poly(I:C), which reportedly binds to the NACHT-LRR region of NLRP1 (*10*), did not affect NLRP1’s interaction with either DPP9 or TRX1 (Fig. S2B,D).

We next wanted to determine if TRX1 binding either restrained or potentiated NLRP1 inflammasome formation. The free NLRP1^CT^ oligomerizes with the adapter protein ASC (apoptosis-associated speck-like protein containing a CARD) to form foci called specks before recruiting CASP1, and VbP induces the formation of visible ASC specks in the cytosol of HEK 293T cells ectopically expressing NLRP1 and GFP-tagged ASC (*14, 23*). Although TRX1 inactivation imposes a fitness cost on cell lines (Fig. S3) (*24*), we managed to generate TRX1*-* deficient HEK 293T cells using sgRNAs targeting *TXN1* (the gene encoding TRX1) (Fig. S4A). We then transiently transfected control (Cas9) or TRX1*-*deficient HEK 293T cells with constructs encoding NLRP1 and GFP-tagged ASC, treated cells with DMSO or VbP, and assessed speck formation by fluorescence microscopy (Fig. S4B, C). We found that TRX1*-*deficient cells had significantly higher numbers of basal and VbP-induced specks compared to control cells. Similarly, we found that siRNA-mediated knockdown of *TXN1* in HEK 293T cells resulted in more basal and VbP-induced NLRP1 inflammasome activation, as evidenced by GSDMD and CASP1 cleavage (Fig. S4D). These data indicated that TRX1 binding restrains the NLRP1 inflammasome.

VbP induces NLRP1-dependent pyroptosis in human immortalized N/TERT-1 keratinocytes (*14*). To evaluate the impact of TRX1 inactivation on NLRP1 inflammasome activation in a physiologically-relevant setting, we next generated TRX1-, NLRP1-, and CASP1-deficient N/TERT-1 keratinocytes using CRISPR/Cas9-based ribonucleoprotein particles (RNPs) (Fig. 3, Fig. S4E,F). We then treated these keratinocytes with VbP for 6 or 18 h before monitoring for signs of pyroptosis, including lactate dehydrogenase (LDH) release, IL-1β/18 cleavage and release, and GSDMD cleavage. Although TRX1 deficiency alone did not cause overt pyroptosis in these cells, VbP induced significant release of LDH and IL-1β/18 only in TRX1-deficient, but not in control, keratinocytes after 6 h (Fig. 4). Thus, TRX1 deficiency accelerated VbP-induced pyroptosis. Moreover, VbP-induced pyroptosis was apparent in both cell types after 16 h, but TRX1-deficient cells nevertheless had released considerably more activated cytokines (Fig. 3). In contrast and as expected, NLRP1- and CASP1-deficient N/TERT-1 keratinocytes were resistant to VbP-induced pyroptosis (Fig. S4F). Overall, these data show that TRX1 binding restrains NLRP1 inflammasome activation.

**Fig. 3.**
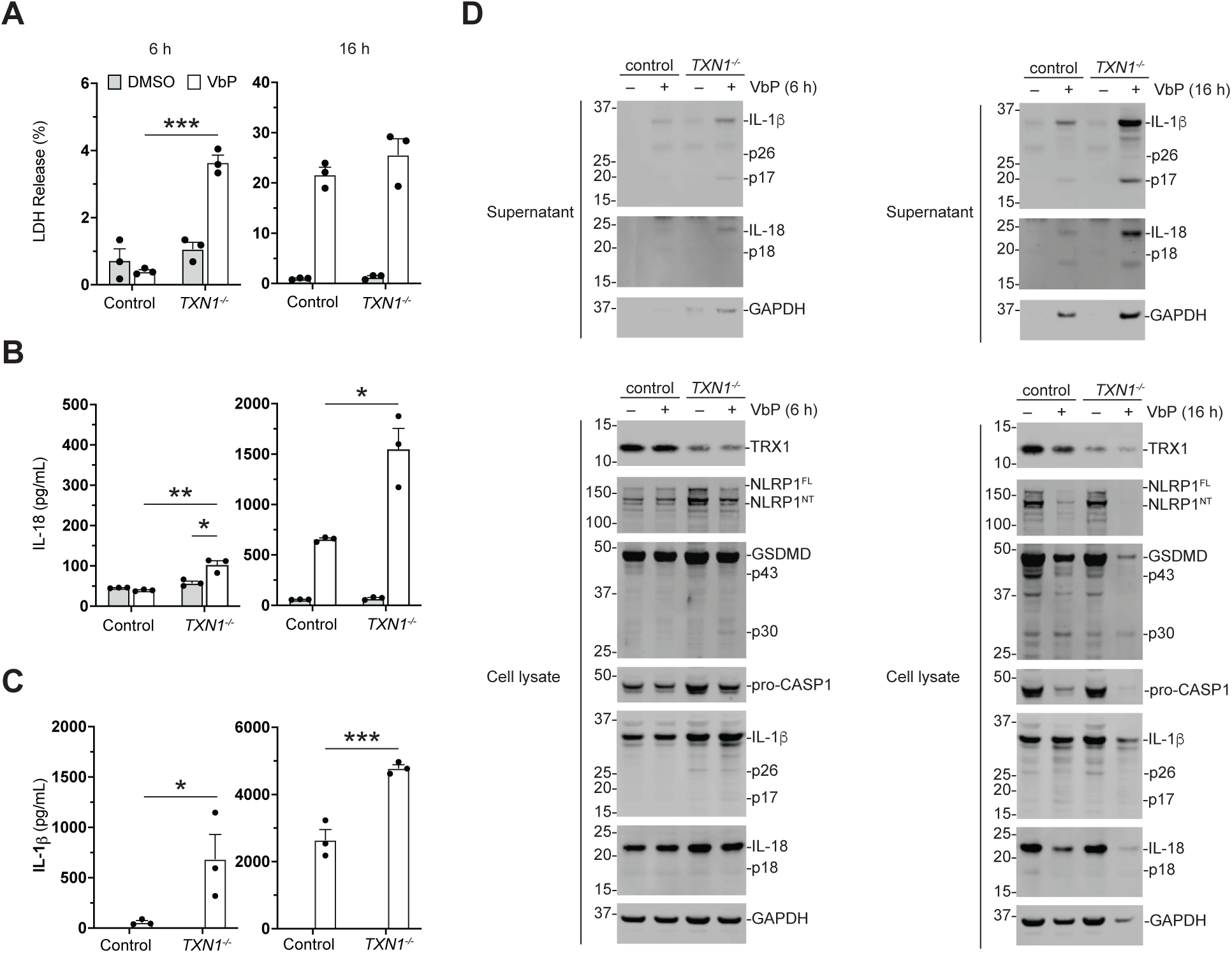
Thioredoxin restrains NLRP1 activation in human keratinocytes. (**A-D**) N/TERT-1 keratinocytes were electroporated with control or Cas9-*TXN1 sgRNA* ribonucleoproteins before being treated with VbP (10 µM) for 6 h or 16 h. Supernatants were analyzed for levels of LDH (**A**), IL-18 (**B**), IL-1β (**C**) release, and both lysates and supernatants were subjected to immunoblot analyses (**D**). Data in **A**-**C** are mean values ± SEM of three biological replicates. * p < 0.05, * * p < 0.01,* * * p < 0.001 as determined by two-sided Student’s *t*-test.

**Fig. 4.**
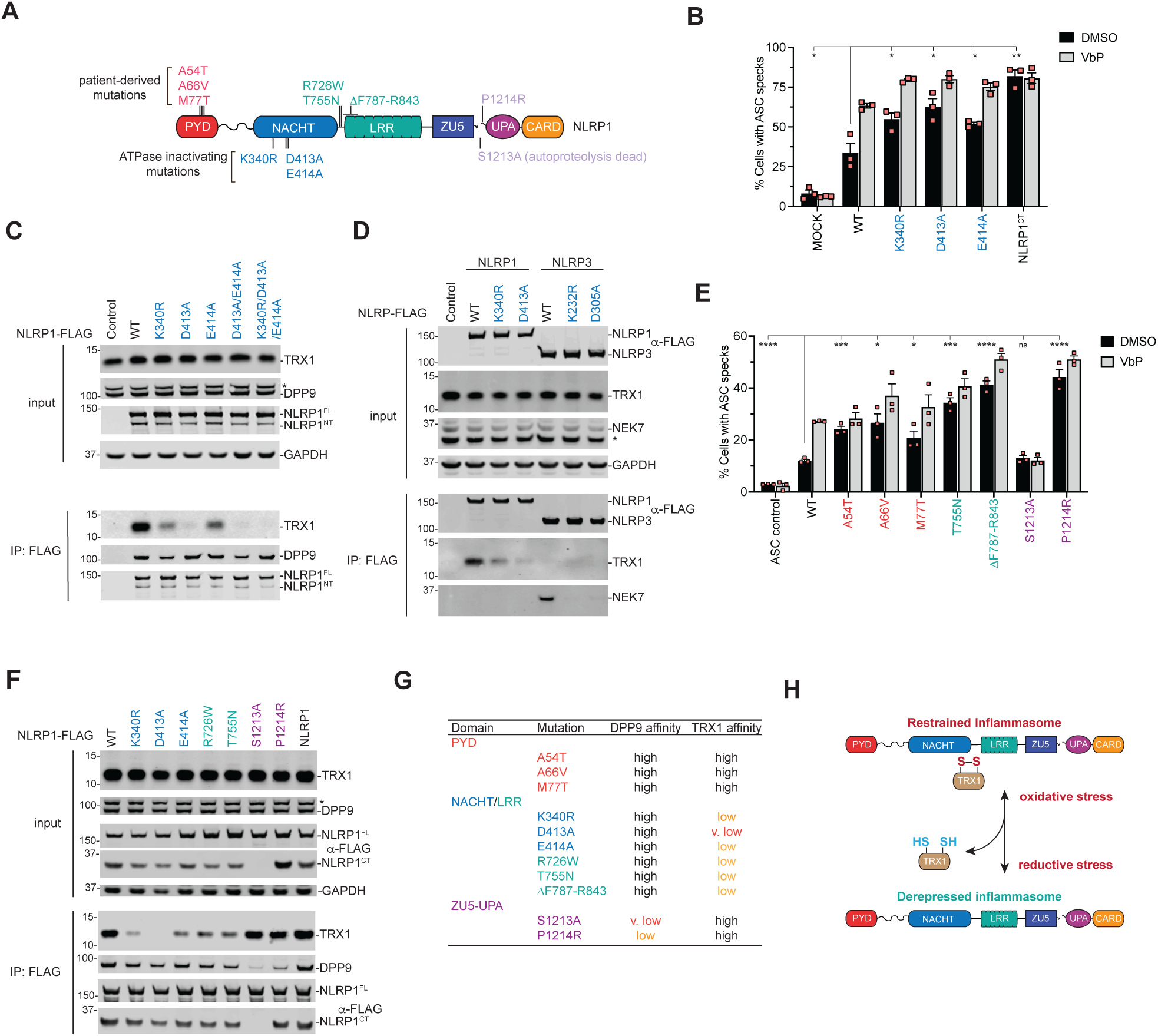
Mutations that abrogate the NLRP1-TRX1 interaction cause inflammasome hyperactivation. (**A**) Diagram depicting the *NLRP1* mutants analyzed. (**B**) HEK 293T cells were transiently transfected with constructs encoding GFP-tagged ASC and the indicated NLRP1 protein, treated with DMSO or VbP (10 µM, 6 h), and evaluated for ASC speck formation by fluorescence microscopy. (**C**-**D**). HEK 293T cells were transfected with the indicated FLAG-tagged NLRP1 or NLRP3 constructs before lysates were harvested, subjected to anti-FLAG IP, and immunoblotted. (**E**) HEK 293T cells were GFP-tagged ASC and the indicated NLRP1 protein and treated and analyzed as in **B**. (**F)**. HEK 293T cells were transfected with the indicated FLAG-tagged NLRP1 or NLRP3 constructs before lysates were harvested, subjected to anti-FLAG IP, and immunoblotted. (**G**) Summary of the impact NLRP1 mutations have on DPP9 and TRX1 binding. (**H)**. Model for TRX1-mediated regulation of the NLRP1 inflammasome. In **B** and **E**, data are mean values ± SEM of three biological replicates. * p < 0.05, * * p < 0.01, * * * p < 0.001 as determined by a two-sided Student’s *t*-test. An asterisk (*) denotes a background band.

The NACHT domains of NLR proteins belong to the signal transduction ATPases with numerous domains (STAND) superfamily of AAA+ ATPases (*25*). These domains use two conserved sequences called Walker A and Walker B motifs to bind and hydrolyze ATP, respectively (*26-28*). Intriguingly, mutation of the Walker A site in NLRP3 inactivates the protein (*26*), but conversely mutation of the Walker A site in mouse NLRP1B generates a hyperactive protein (*29, 30*). The mechanistic basis for this striking difference is not yet understood. Consistent with these previous reports, we found that mutation of either the Walker A (K340R) or B (D413A or E414A) sites in human NLRP1 similarly causes spontaneous NLRP1 inflammasome activation in the HEK 293T ASC speck assay (Fig. 4A,B, Fig. S5A,B). These mutant proteins, unlike the free NLRP1^CT^, were still restrained at least in part by DPP9, as VbP still induced additional inflammasome formation (Fig. 4B, Fig. S5A).

We next hypothesized that ATPase activity might be required for NLRP1 to form the repressive complex with TRX1. Indeed, we found that mutation of either the Walker A or Walker B site weakened the NLRP1-TRX1 interaction (Fig. 4C,D), and mutation of both sites (K340R/D413A/E414A) completed abrogated the interaction. Notably, these mutant proteins still associated with DPP9 (Fig. 4C). Intriguingly, we analogously found that mutation of the Walker A (K232R) or B (D305A) site in NLRP3 disrupted its interaction with NEK7 (Fig. 4D). Thus, the NACHT-LRR regions of both NLRP1 and NLRP3 require ATP binding and hydrolysis to associate with their endogenous binding partners. However, NLRP1 forms a complex with TRX1 that represses inflammasome activation, whereas NLRP3 forms a complex with NEK7 that licenses inflammasome activation. Overall, these data explain the contrasting requirements of ATPase activity in the NLRP1 and NLRP3 inflammasomes.

A number of germline-encoded mutations in *NLRP1*, including several in the NACHT-LRR region, cause autoinflammatory disorders (Fig. 4A) (*31-34*). We confirmed that patient-derived mutations in the PYD, NACHT-LRR, and FIIND domains promoted spontaneous inflammasome assembly relative to wild-type (WT) NLRP1 in the HEK 293T ASC speck assay (Fig. 4E, Fig. S5C,D). The P1214R mutation, which occurs within the NLRP1 FIIND, is hyperactive due to diminished DPP9 binding (*6, 14*), and we reasoned that the NACHT-LRR mutations might analogously dysregulate the inflammasome by impairing the interaction with TRX1. Indeed, we found that the three patient-derived mutations (R726W, T755N, and ΔF787-R843) in the NACHT-LRR region of human NLRP1 (Fig. 4F, Fig. S6A-C) reduced binding to TRX1 to a similar extent as the Walker A (K340R) mutation (Fig. 4C, Fig. S6C). Similarly, a Q593P mutation in the NACHT-LRR region of the mouse NLRP1A, which causes neutrophilic inflammatory disease in mice (*34*), weakened its association with TRX1 (Fig. S6D). In contrast to the NACHT-LRR mutations, mutations in the PYD (A54T, A66V, M77T) and FIIND (P1214R) domains did not impact the TRX1-NLRP1 interaction (Fig. 4F, Fig. S6C,E). As expected, only the patient-derived P1214R FIIND mutant underwent full autoproteolysis and had impaired binding to DPP9 (Fig. 4F, Fig. S6C). It should be noted that several mutations, in particular the R726W mutation in human NLRP1 and the Q593P mutation in mouse NLRP1A, appeared to weaken DPP9 binding (Fig. S6C,D), but this is likely due to compromised autoproteolysis. Thus, patient-derived mutations in NACHT-LRR and FIIND cause inflammatory disorders by disrupting the repressive interactions with TRX1 and DPP9, respectively (Fig. 4G). The PYD mutations are thought to directly destabilize the domain’s fold (*31*), possibly increasing the susceptibility of the NLRP1^NT^ to proteasome-mediated degradation (*35*).

Although only ∼5-10% of TRX1 is oxidized in the cytosol of unstressed human cells (*36*), TRX1 is among the most abundant proteins in mammalian cells and therefore a sufficient level of oxidized TRX1 should normally exist to repress the much less abundant NLRP1 protein (*37*). Indeed, our genetic knockout and mutation data above shows that TRX1 restrains the NLRP1 inflammasome in cells. We reasoned that lowering the intracellular redox potential (i.e., inducing reductive stress) should decrease the fraction of oxidized TRX1, dissociate TRX1 from NLRP1, and potentiate inflammasome activation. Unfortunately, however, we are not aware of any agent known to lower the intracellular redox potential and decrease the fraction of oxidized TRX1. Regardless, we next wanted to assess the impact of a panel of (anti)oxidants, including DTT, N-acetyl-cysteine (NAC), H2O2, and antimycin, on NLRP1 inflammasome activation (Fig. S7A). Although several of these agents were toxic on their own at high doses, none impacted basal- or VbP-induced pyroptosis in human NLRP1-dependent N/TERT-1 keratinocytes (Fig. S7B,C), mouse NLRP1B-dependent RAW 264.7 cells (Fig. S7D-F), or human CARD8-dependent MV4;11 cells (Fig. S7G). Consistent with our results, DTT did not substantially reduce a fluorescent biosensor for TRX1 redox (TrxRFP1) in HEK 293T cells below its basal oxidation state (*38*). Thus, our data strongly indicate reductive stress will potentiate NLRP1 activation, but additional research is needed to identify and characterize reagents that induce reductive stress and alter the TRX1 redox couple in cells.

In summary, we have now established that oxidized, but not reduced, TRX1 restrains NLRP1 inflammasome activation by directly associating with the NLRP1 NACHT-LRR region. This suggests that reductive stress is a danger signal that contributes to NLRP1 inflammasome activation, and conversely that oxidative stress represses its activation (Fig. 4H). Projecting forward, it will be important to determine not only why reductive stress is an inflammasome-activating danger signal, but also how TRX1 binding represses inflammasome formation. On that note, we recently discovered that DPP8/9 inhibition accelerates the degradation of disordered and misfolded proteins through a poorly characterized proteasome pathway (*17*). As such, we hypothesize that TRX1 dissociation might increase the disorder of the NLRP1^NT^ fragment, thereby increasing its susceptibility to degradation by this pathway. Future studies are needed to investigate the relationship between the cellular redox state, disordered protein degradation, and inflammasome activation. Regardless, our study here now reveals a new fundamental mechanism that links the intracellular redox environment to the activation of the innate immune system.

## Acknowledgments

We thank Bachovchin lab for helpful discussions.

## Author contributions

D.P.B., A.E.W., C.D.W., Q.W., A.R.G., and S.D.R. designed experiments, performed experiments, and analyzed data. D.A.B. directed the project, designed experiments, and analyzed data. D.P.B. and D.A.B. wrote the paper.

## Data and materials availability

All data are available in the main text or the supplementary materials.

## Supplementary Materials

Materials and Methods

Figures S1-S7

Tables S1

References *40-42*)

---

**Fig. S1.**
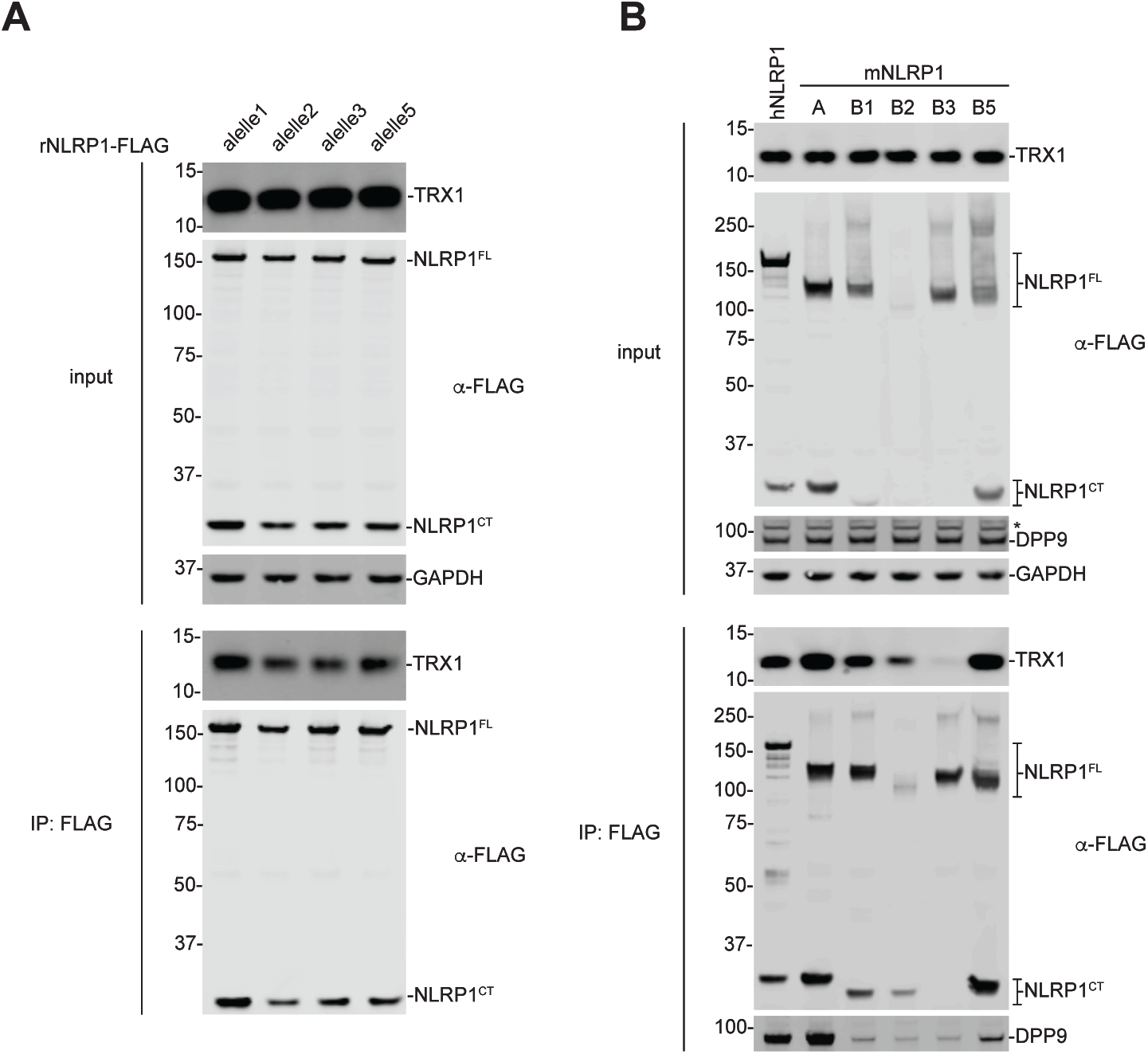
The functional rodent NLRP1 proteins bind TRX1. (**A**,**B**) HEK 293T cells were transiently transfected with the indicated FLAG-tagged rat NLRP1 (rNLRP1) alleles (**A**) or human (hNLRP1) or mouse NLRP1 (mNLRP1) alleles (**B**), subjected to anti-FLAG IP, and analyzed by immunoblotting. It should be noted that NLRP1B3 is non-functional, and, unlike the other rodent alleles, neither binds to TRX1 nor undergoes autoproteolysis.

**Fig. S2.**
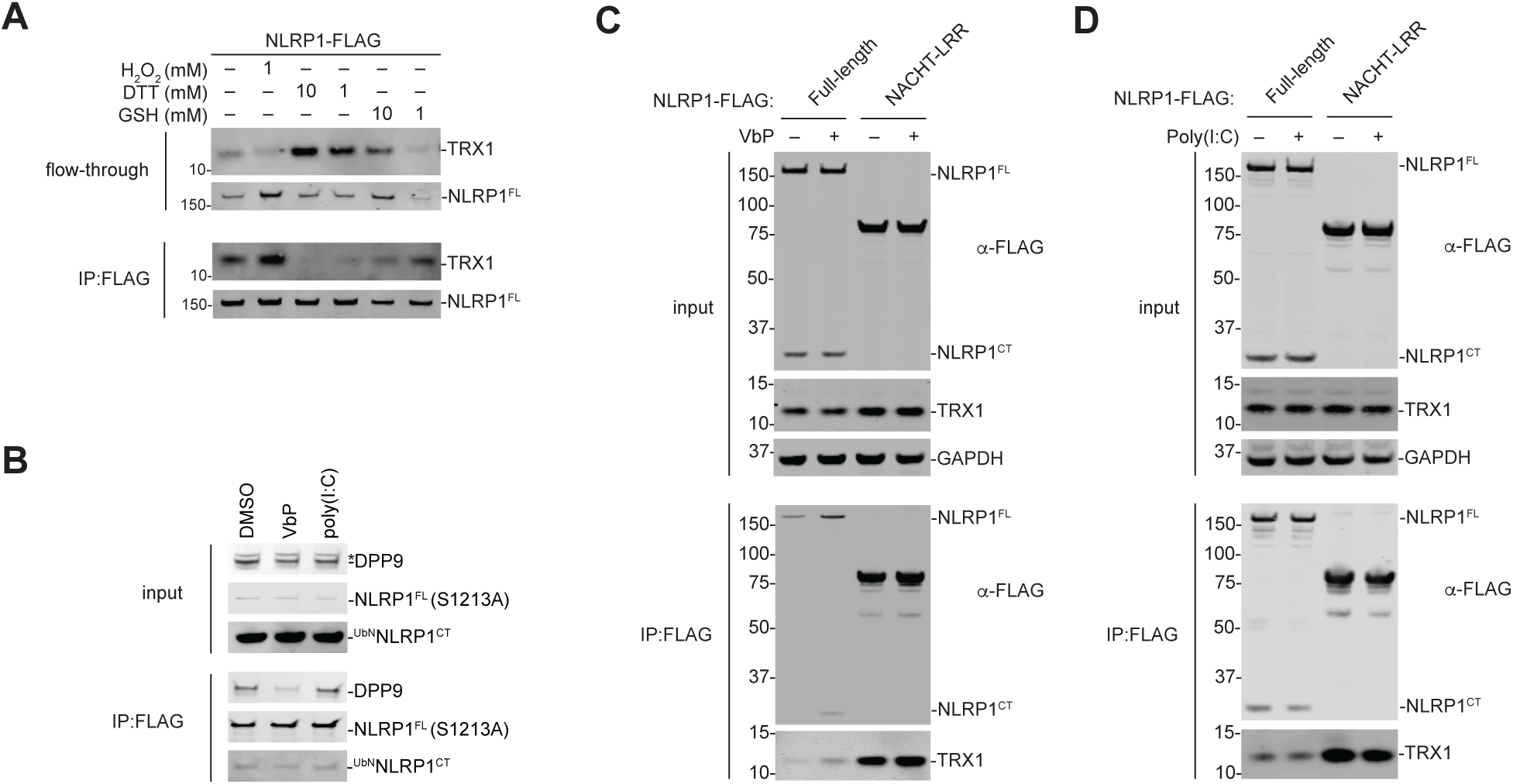
Antioxidants, but not poly(I:C) or VbP, disrupt the NLRP1-TRX1 interaction. (**A-D**) HEK 293T cells were transfected with the indicated constructs before lysates were harvested and subjected to anti-FLAG IP and immunoblotting analysis. Anti-FLAG beads were treated with (anti)oxidants (**A**), VbP (10 µM, **B, C**) or high molecular weight poly(I:C) (3.33 µg/mL, **B**,**D**). The flow-through fraction in **A** is the unbound fraction after (anti)oxidant treatment.

**Fig. S3.**
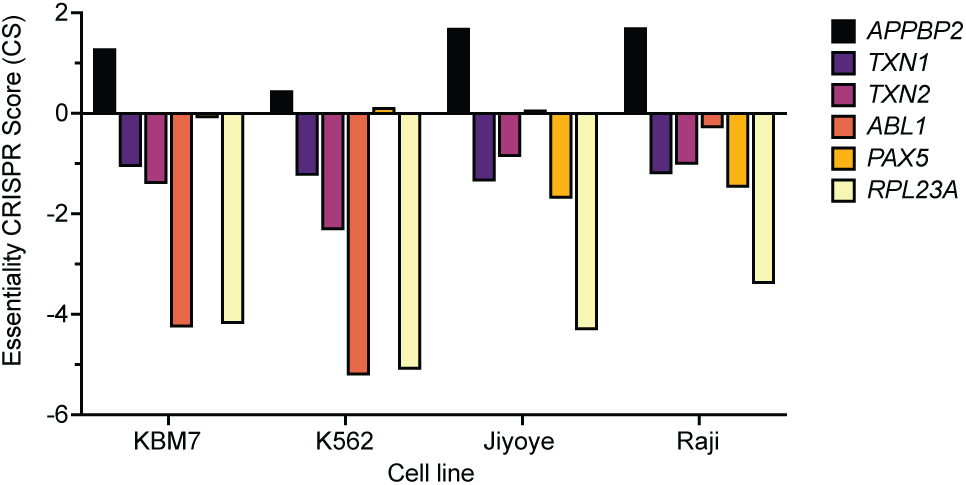
*TXN* knockout compromises cell viability. Gene essentiality scores were obtained from a manuscript that used genome-wide sgRNA screens to identify genes required for proliferation and survival (*42*).

**Fig. S4.**
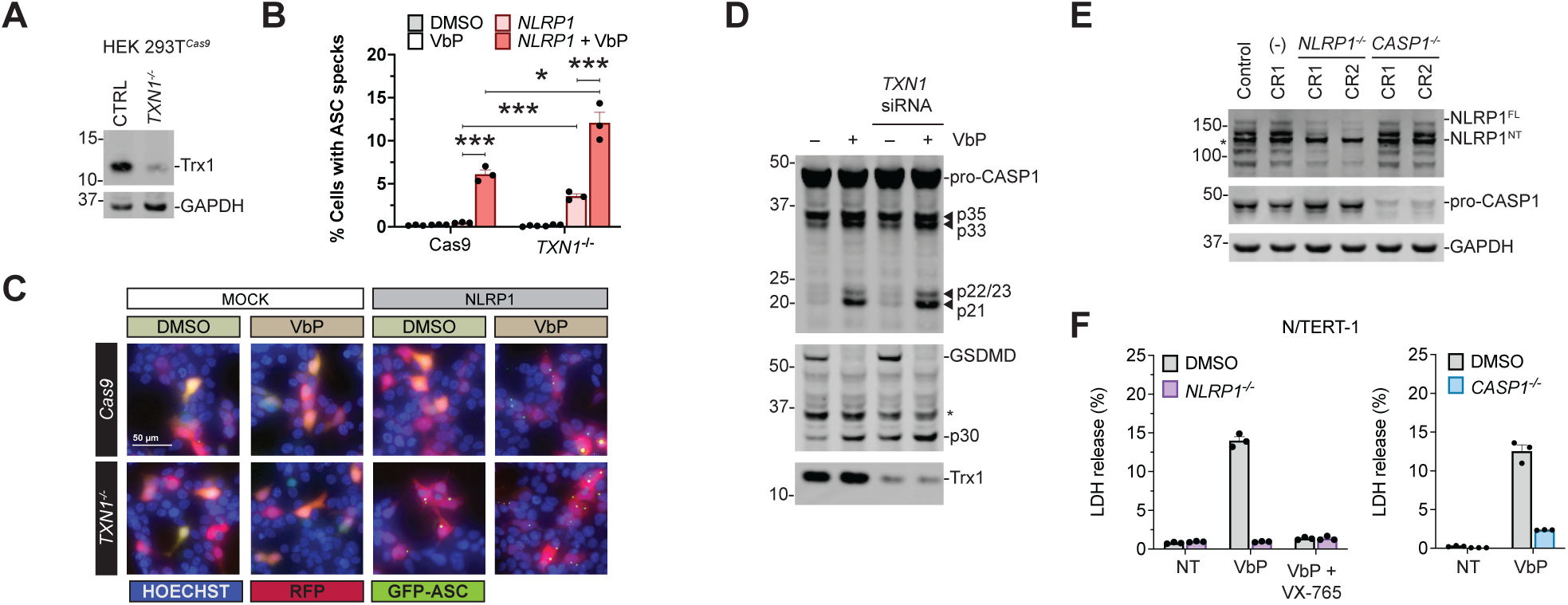
Genetic loss of *TRX1* causes NLRP1 inflammasome hyperactivation. (**A**) sgRNAs targeting *TXN1* were used to generate TRX1-deficient HEK 293T cells. Loss of TRX1 was confirmed by immunoblotting. (**B**-**C**) Control (Cas9) and *TXN1*^*-/-*^ HEK 293T were transiently transfected with constructs encoding GFP-tagged ASC and NLRP1, treated with DMSO or VbP (10 µM, 6 h), and evaluated for ASC speck formation by fluorescence microscopy. (**D**) HEK 293T cells stably expressing CASP1 and GSDMD were treated with an siRNA targeting *TXN1* before transfection with constructs encoding ASC and NLRP1. Cells were then stimulated with VbP (10 µM, 6 h) and analyzed by immunoblotting. (**E**) RNPs containing the indicated *sgRNA*s were used to generate knockout N/TERT-1 cells. Protein loss was confirmed by immunoblotting. (**F**) Control, *NLRP1*^-/-^, and *CASP1*^*-/-*^ N/TERT-1 keratinocytes were treated DMSO or VbP (10 µM) for 16 h before LDH release was evaluated. The indicated samples were pretreated with the CASP1 inhibitor VX-765 (10 µM) for 30 min before the addition of VbP. In **B** and **F**, data are mean values ± SEM of three biological replicates. * p < 0.05, * * p < 0.01, * * * p < 0.001 as determined by a two-sided Student’s *t*-test. An asterisk (*) denotes a background band.

**Fig. S5.**
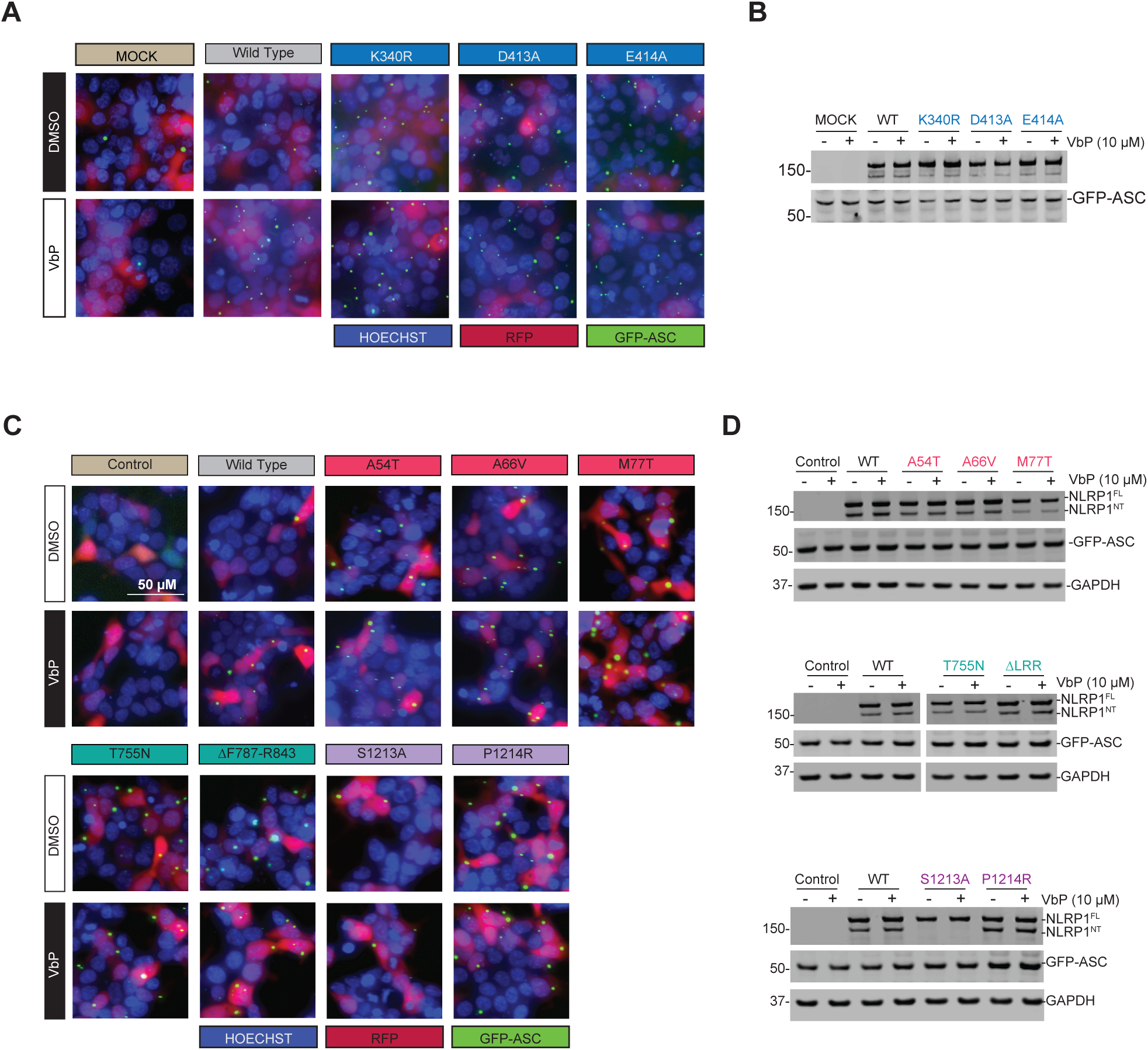
ATPase-inactivating and patient-derived mutations hyperactivate NLRP1. (**A-D**) HEK 293T cells were transiently transfected with constructs encoding GFP-tagged ASC and the indicated NLRP1 protein, treated with DMSO or VbP (10 µM, 6 h), and evaluated for ASC speck formation by fluorescence microscopy. Representative images of GFP-ASC foci formation are shown in **A** and **C**. Protein expression was confirmed by immunoblotting in **B** and **D**.

**Fig. S6.**
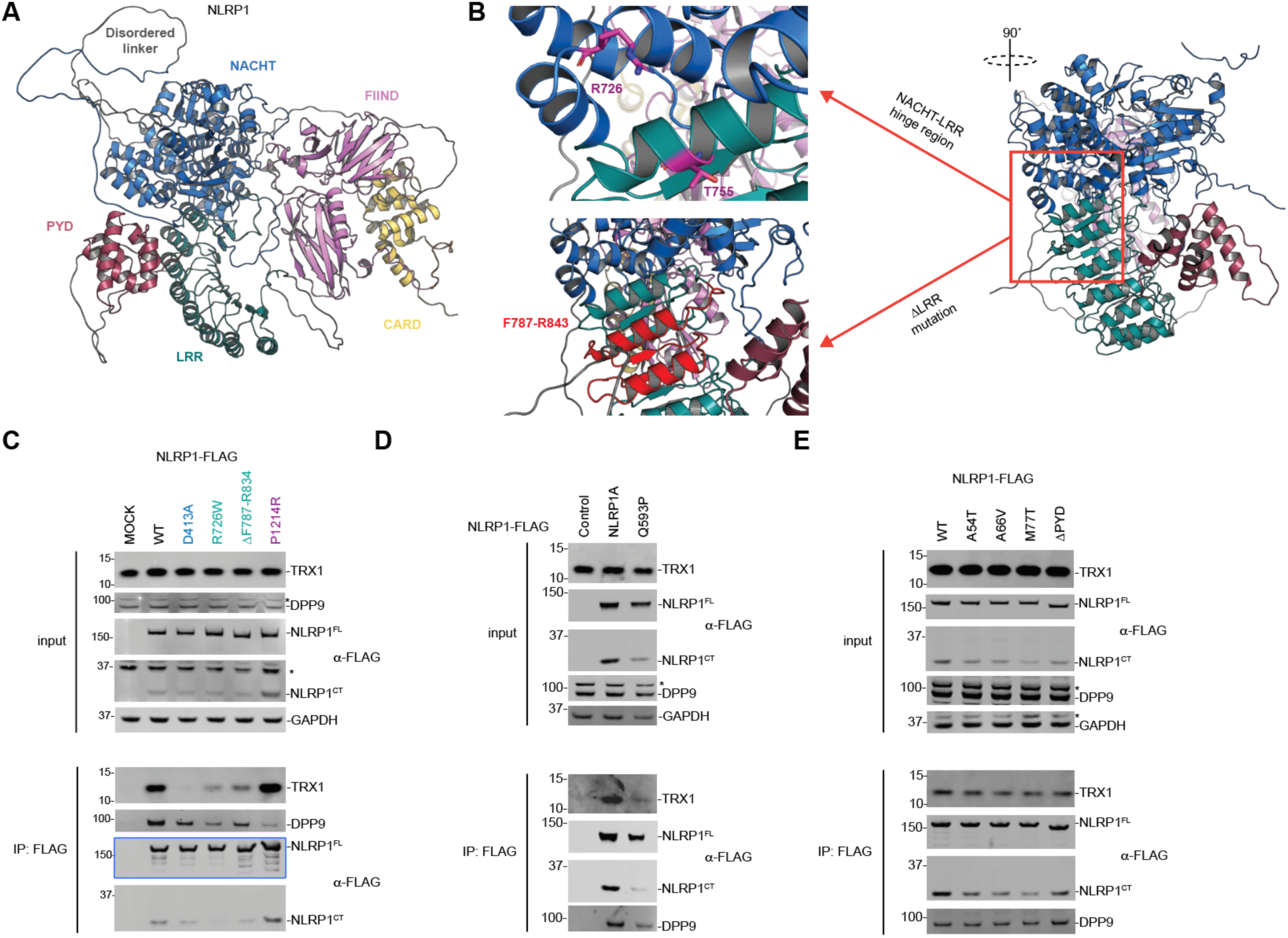
Mutations within the NACHT-LRR domain of NLRP1 weaken TRX1 binding. (**A**) Structure prediction of human NLRP1 by AlphaFold Protein Structure Database (*43*). (**B**) Predicted locations of R726W and T755N (left, top) and Δ F787-R843 (left, bottom) mutations at the NACHT-LRR interface. (**C**-**E**) HEK 293T cells were transfected with constructs encoding the indicated FLAG-tagged NLRP1 constructs before lysates were harvested, subjected to anti-FLAG IP, and immunoblotted.

**Fig. S7.**
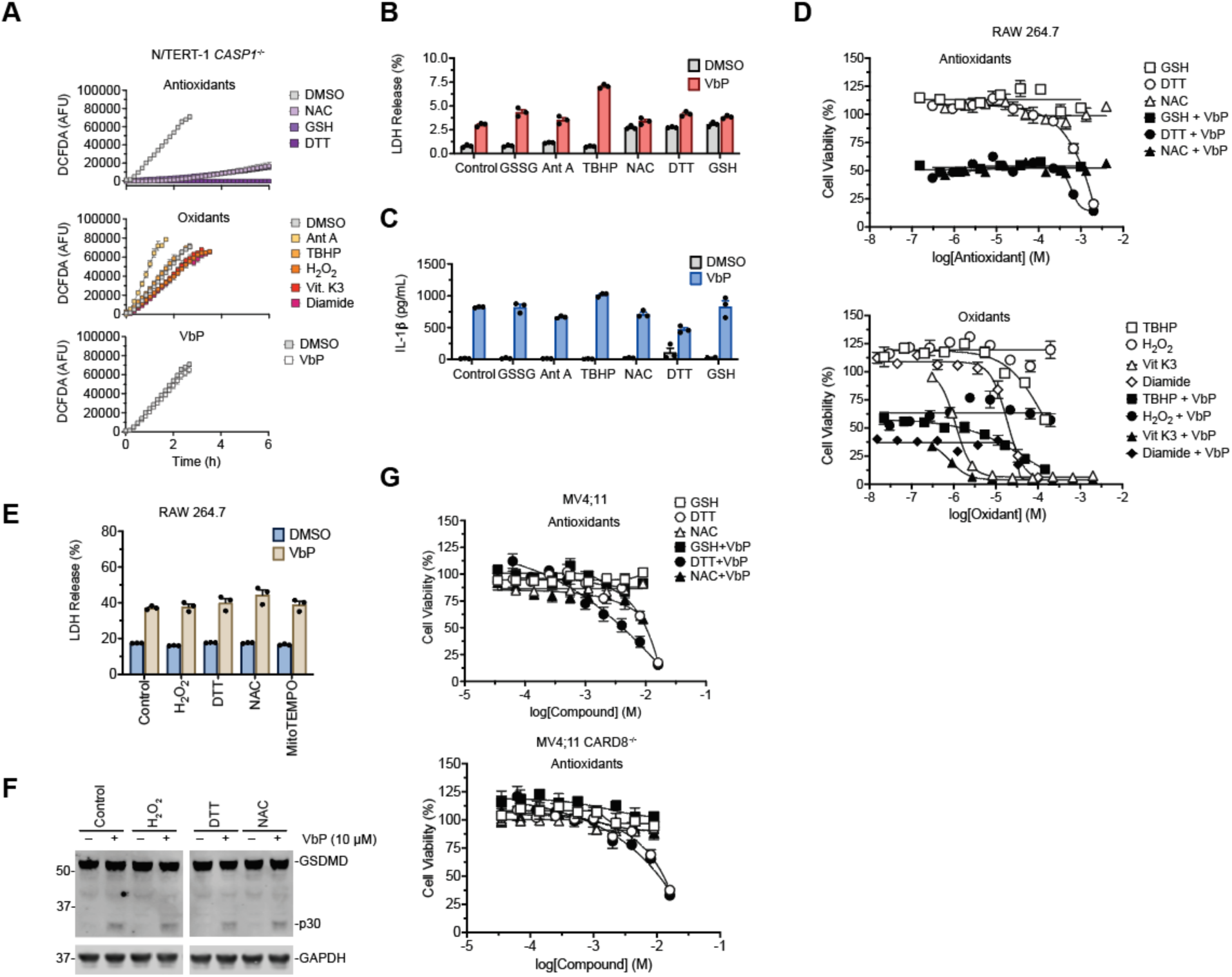
(Anti)oxidant treatment does not impact basal or VbP-induced CARD8 or NLRP1 activation. (**A**) Cellular ROS levels were measured in *CASP1*^*-/-*^ N/TERT-1 keratinocytes (used to prevent pyroptosis) following treatment with the indicated (anti)oxidants using the cell permeable 2’,7’-dichlorodihydrofluorescein diacetate dye (DCFDA), which fluoresces upon oxidation. (**B**) Wild type N/TERT-1 keratinocytes treated with the indicated (anti)oxidants alone or together with VbP (10 µM) for 6 h before LDH and IL-1β release into supernatants was evaluated. (**D**-**F**) RAW 264.7 cells were treated with varying doses of the indicated (anti)oxidants alone or with VbP (10 µM) for 6 h before cell viability was assessed by Cell-TiterGlo (CTG, **D**) and by LDH release assay (**E**) and immunoblot (**F**) for single dose (anti)oxidant concentration treatment. (**G**) MV4;11 wild type (top) and *CARD8*^*-/-*^ (bottom) cells were treated with varying doses of the indicated (anti)oxidants alone or with VbP (10 µM) for 6 h before cell viability was assessed by Cell-TiterGlo (CTG). Unless otherwise indicated, GSSG, GSH, DTT, NAC, TBHP, and H2O2 are treated at 100 µM, while Vit. K3, Antimycin A, VbP, and MitoTEMPO are treated at 10 µM, and Diamide was treated at 5 µM. All statistical data are means ± SEM for at least 3 biological replicates. * p < 0.05, * * p < 0.01, * * * p < 0.001.

**Table S1.** Potential NLRP1 binding proteins identified by IP-MS analyses. Proteins identified by mass spectrometry in the NLRP1-FLAG IP experiment were ranked in descending order of abundance ratio. This data table associated with main text Fig. 1B.

| Rank | Protein | NLRP1-FLAG<br>Abundance | GFP-FLAG<br>Abundance | Abundance Ratio<br>(NLRP1/GFP) | Adj. p-value |
| --- | --- | --- | --- | --- | --- |
| 1 | **DPP8** | 178.7 | 18.9 | 9.444 | 0.104 |
| 2 | **NLRP1** | 118095 | 13179.2 | 8.961 | 0.025 |
| 3 | **DPP9** | 9879.8 | 1140.2 | 8.665 | 0.084 |
| 4 | ARL11 | 159.6 | 19 | 8.402 | 0.053 |
| 5 | ATP10A | 188.2 | 28 | 6.729 | 0.045 |
| 6 | **TRX1** | 15979.7 | 2561.6 | 6.238 | 0.044 |
| 7 | PPM2C | 232.5 | 40.7 | 5.706 | 0.159 |
| 8 | TUBA1A | 4707.1 | 844 | 5.577 | 0.087 |
| 9 | FLJ77553 | 804.4 | 145.1 | 5.544 | 0.241 |
| 10 | MIC13 | 152.1 | 27.5 | 5.536 | 0.147 |
| 11 | DNAJA1 | 8681.2 | 1579.9 | 5.495 | 0.036 |
| 12 | TUBA4A | 1235.9 | 249.4 | 4.956 | 0.044 |
| 13 | IFT81 | 64.4 | 13.4 | 4.799 |  |
| 14 | FLJ12755 | 88.4 | 18.9 | 4.681 | 0.086 |
| 15 | FBXW8 | 81.4 | 17.8 | 4.566 | 0.125 |
| 16 | ANP32B | 6258.5 | 1400.3 | 4.469 | 0.063 |
| 17 | TUBB4A | 588.8 | 132.9 | 4.431 | 0.044 |
| 18 | TUBA1C | 37789.2 | 8553.7 | 4.418 | 3.89E-13 |
| 19 | SET | 3513.8 | 857.5 | 4.098 | 0.053 |
| 20 | TUBB4B | 6065.7 | 1498.3 | 4.048 | 0.013 |

## Materials and Methods

### Plasmids, antibodies, and reagents

*NLRP1* and *CARD8* transcript variant 1 DNA obtained commercially from Origene (pCMV6-entry clones RC216481 and RC230245, respectively). *NLRP3, TXN1, Nlrp1a, and Nlrp1b2* DNA was purchased from Genscript (OHu26815D, OHu15866D, OMu19634, and OMu00866D, respectively). DNA encoding *Nlrp1b1* was cloned from RAW 264.7 macrophages. DNA encoding the mouse *Nlrp1b3* and *Nlrp1b5* were obtained from R. Vance and J. Mogridge. DNA encoding the rat *Nlrp1* alleles 1-5 were cloned from Sprague Dawley, CDF, Zucker, Copenhagen, and Lewis rodents, respectively. Expression constructs were first amplified by polymerase chain reaction (PCR) and subcloned into pDONR221, then into modified pLEX307 vectors originating from pLEX307 (Addgene #41392) using Gateway cloning technology (ThermoFisher Scientific). *TXN1* CRISPR *sgRNA*s were cloned into the pLentiGuide expression system (Addgene #52963) for HEK 293T gene ablation. Viral transductions were performed with the psPAX2 lentiviral packaging vector (Addgene #11260) and VSV-G envelope expression plasmid, pMD2.G (Addgene #12259). Unless otherwise indicated, all plasmid transfections were performed with FuGene HD (Promega) following manufacturers recommendations. For siRNA protein knockdown studies, unless otherwise indicated, targeting or non-targeting sequences were transfected into cells with Lipofectamine RNAiMAX (ThermoFisher Scientific) following the manufacturer’s recomendations. For Cas9 Ribonucleoprotein (RNP) complex genetic modifications, the Alt-R CRISPR-Cas9 system was used with pre-designed crRNA for *NLRP1, CASP1, TXN1, and with* tracrRNA and Alt-R S.p. Cas9 Nuclease V3 recombinant protein (Integrated DNA Technologies, IDT), which were delivered to cells using the Neon transfection system (ThermoFisher Scientific). Point mutations were prepared using QuikChange II site-directed mutagenesis (Agilent), following manufacturers suggested protocols. Antibodies used in this study were: anti-FLAG antibody (F1804, Millipore SiGMa), anti-NEK7 (3057S, Cell Signaling Technology), anti-TRX1 (ab86255, Abcam; 2429S, Cell Signaling Technology), anti-DPP9 (ab42080, Abcam), anti-GAPDH (2118S, Cell Signaling Technology), anti-NLRP1 (AF6788, R&D systems), anti-CARD8 (ab194585, Abcam), anti-ASC (AF3806, R&D systems), anti-CASP1 (2225S, Cell Signaling Technology; ab179515, Abcam), anti-GSDMD (NBP2-33422, Novus), anti-mGSDMD (ab209845, Abcam) anti-IL-1β (AF201NA, R&D systems), anti-IL-18 (AF2548, R&D systems).

Experiments involving Poly(I:C) were performed using the long synthetic analog of dsRNA, Poly(I:C) HMW (InvivoGen). Transfection of Poly(I:C) was performed using Lipofectamine 2000 (ThermoFisher Scientific) following manufacturers recommendations. VX-765 was obtained from APExBio; Antimycin A from *Streptomyces sp*., *N*-acetyl cysteine (NAC), *DL-*dithiothreitol, hydrogen peroxide, Diamide, and MitoTEMPO were obtained from Sigma Aldrich; *tert*-butyl hydrogen peroxide was obtained from ACROS Organics; Oxidized and reduced *L*-Glutathione were obtained from Alfa Aesar; Val-boroPro and Vitamin K3 were obtained from Cayman Chemical Company; HOECHST 33342, trihydrochloride, trihydrate was obtained from Invitrogen.

### Cell culture

HEK293T and RAW264.7 were purchased from ATCC and cultured in Dulbecco’s Modified Eagle’s Medium (DMEM) with *L*-glutamine and supplemented with 10% fetal bovine serum (FBS) and 1X penicillin/streptomycin. MV4;11 cells were purchased from the Liebnez Institute DSMZ’s German collection of microorganisms and cell cultures GmbH and cultured in Roswell Park Memorial Institute (RPMI) medium 1640 with *L*-glutamine supplemented with 10% FBS and 1X penicillin/streptomycin. N/TERT-1 cells, a gift from James Rheinwald (*40*), were cultured in Keratinocyte serum free medium (KSFM, Gibco) supplemented with 100 U/mL penicillin (Gibco), 100 µg/mL streptomycin (Gibco), bovine pituitary extract (20 µg/ml, Gibco) and recombinant epidermal growth factor (EGF, 2 ng/ml, Gibco).

All cells were grown at 37 °C in a 5% CO2 atmosphere incubator. Cell lines were regularly tested for mycoplasma using the MycoAlert™ Mycoplasma Detection Kit (Lonza).

### TMT labeling for IP-MS/MS analysis

3×10^6^ HEK 293T cells were seeded in a 10 cm tissue culture dish and transfected the following day with an expression plasmid for the given FLAG-tagged protein construct. The cells were harvested 48 h post transfection and lysed in 1 mL of 0.5% NP-40 on ice for 30 min. Lysate samples were placed in a tabletop centrifuge for rotation at 2000 x *g* for 10 min at 4 °C. The soluble fraction was separated, and protein content normalized (DCA protein assay, BIO-RAD). Protein samples were globally reduced by application of 10 mM TCEP (tris(2-carboxyethyl) phosphine) for 1h at 56 °C with shaking, followed by alkylation of free thiols with 20 mM iodoacetamide for 30 min at ambient temperature, protected from light. Protein samples were then precipitated from solution by adding -20 °C acetone to six sample volumes and incubated overnight at -20 °C. Precipitated protein was isolated by sample rotation at 8000 x *g* for 10 min at 4 °C and decanting the liquid phase. The pellet samples were resuspended in in 100 µL 50 mM triethylammonium bicarbonate (TEAB) followed by total protein quantification by DCA assay prior to digestion with 2.5 mg trypsin per 100 mg sample at 37 °C overnight.

TMTsixplex sample reagents (ThermoFisher Scientific) were equilibrated to ambient temperature and 0.8 mg of label (126, 127 = GFP-FLAG; 128, 129 = NLRP1-FLAG; 130, 131 = unused sample) diluted into 41 µL pure acetonitrile before use. 41 µL of each TMT label reagent were added to each protein sample and incubated at ambient temperature for 1 h. 8 µL of a 5% hydroxylamine solution was added to quench each sample and incubated at ambient temperature for 15 min. Samples were then equally combined and purified by a High pH Reversed-Phase Peptide Fractionation Kit (Pierce) and dried with a Genevac EZ-2 evaporator to yield peptide pellets submitted for tandem LC-MS/MS/MS analysis.

### TMT LC-MS/MS/MS

Mass spectrometry data was collected on an Orbitrap Fusion Lumos mass spectrometer coupled to an Easy-nLC 1200 (ThermoFisher Scientific). Peptides were separated over a 200 min gradient of 0-50% acetonitrile in water with 0.1 % formic acid at a flow of 300 nL/min on a 50 cm PepMap RSLC C18 column (2 mm, 100 Å, 75 µm x 50 cm). The full MS spectrum was acquired in the Orbitrap at a resolution of 120,000. The 10 most intenst MS1 ions were selected for MS2 analysis. The isolation width was set at 0.7 m/z and isolated precursors were fragmented by CID (35% CE). Following acquisition of each MS2 spectrum, a synchronous precursor selection (SPS) MS3 scan was collected on the top 10 most intense ions in the MS2 spectrum. The isolation width was set at 1.2 m/z and isolated precursors were garmented using HCD. The mass spectrometry proteomics data have been submitted to the ProteomeXchange Consortium (http://proteomecentral.proteomexchange.org) via the PRIDE partner repository (*41*).

### Proteomic analysis

MS raw files were converted to MGF and processed using Proteome Discoverer version 2.2.0.388 (Thermo Scientific) by searching against the Uniprot human database supplemented with common contaminant protein sequences and quantified according to SPS MS3 reporter ions. Identification was based on CID spectra using SequestHT. Search criteria included: 20 ppm mass tolerance for MS spectra, 0.5 Da fragment mass tolerance for MS/MS spectra, a maximum of two allowed missed cleavages, static carbamidomethylation (+57.021 Da) of cysteine and TMTsixplex (+229.163 Da) of lysine and of peptide N-terminus, dynamic modifications of methionine oxidation (+15.995 Da), N-terminal protein acetylation (+42.011 Da), asparagine or glutamine deamidation (+0.984 Da), and serine, tyrosine, or tryptophan phosphorylation (+79.966 Da) and a false discovery rate of < 0.01.

### Knockout cell lines

*TXN1*^*-/-*^ HEK 293T cells were prepared by transient transfection of HEK 293T cells stably expressing *Cas9* with *sgRNA* for *TXN1* packaged in lentiGuide-Puro (Addgene, 52963) using the following oligo sequences (5’-ACGTGATATTCCTTGAAGTA-3’; 5’-TAGTTGACTTCTCAGCCACG-3’, 5’-GGTGAAGCAGATCGAGAGCA-3’). 48 h following transfection, cells were selected with 1 µg/mL puromycin for 5 days and subjected to single cell cloning protocols to obtain a population enriched for the gene modification.

*CASP1*^*-/-*^ and *NLRP1*^*-/-*^ N/TERT-1 keratinocyte cells were prepared by delivery of Cas9 ribonucleoprotein complexes containing an Alt-R CRISPR-Cas9 sgRNA and recombinant Cas9 (IDT) using the Neon Transfection System (ThermoFisher Scientific) following the manufacturer’s recommendations. Briefly, sgRNA complexes were prepared by combining predesigned Alt-R CRISPR-Cas9 crRNA (*NLRP1*: 5’-GGTGGTAGGAACGCCCCCAC-3’; CR2, 5’-CTGGATCCATGAATTGCCGG-3’. *CASP1*: CR1, 5’-CGGCTTGACTTGTCCATTAT-3’; CR2, 5’-GACCTCTGACAGCACGTTCC) with Alt-R CRISPR-Cas9 tracrRNA to 44 µM and annealing by heating to 95 °C for 5 min followed by gradual cooling to ambient temperature over 30 min. To form the RNP complexes sgRNA samples and recombinant Alt-R Cas9 enzyme were combined and incubated for 20 min. RNP complexes were then introduced to 5×10^5^ wild type N/TERT-1 keratinocyte cells with Alt-R Cas9 Electroporation Enhancer (IDT) and electroporated with 2 pulses of 1700 V for 10 ms. Cells were then transferred to warm supplemented KSFM without antibiotics to recover for 72 h, expanded, and evaluated by immunoblot for protein expression levels prior to experimentation.

To prepare *TXN1*^*-/-*^ N/TERT-1 keratinocytes, 5×10^5^ cells were electroporated using the same protocol as above, with the exception that RNP complexes comprised a mixture of three sgRNAs (CR1, 5’-TAGTTGACTTCTCAGCCACG-3’; CR2, 5’-GACTTCTCAGCCACGTGGTG-3’; CR3, 5’-ACTTCTCAGCCACGTGGTGT-3’) and cells were electroporated as individual replicates prior to each experiment.

### siRNA Knockdown

2.5×10^5^ HEK 293T cells were seeded in 6-well tissue culture plates and incubated overnight to adhere. The following day non-targeting to *TXN1* Silencer pre-designed siRNA (Ambion) were transfected to cells using Lipofectamine RNAiMAX (ThermoFisher Scientific) following the manufacturer’s recommendations.

### Western Blotting

For cell lysate samples, cell pellets were resuspended in 1X PBS and mechanically lysed by sonication before protein standardized by DC assay (BIO-RAD) and mixed with Western blot sample loading buffer (2X loading buffer (LI-COR), 100 mM DTT) before heat-denaturation at 97 °C for 10 min.

For supernatants, protein pellets were prepared by precipitation with either MeOH/CHCl3 extraction or by addition of four sample volumes of acetone at -20 °C followed by centrifugation at 3000 x *g* for 30 min at 4 °C and decanting. Protein pellets were then suspended in 1X PBS and combined 1:1 with 2X sample loading buffer before heating to 97 °C for 10 min.

All protein samples were separated by SDS-PAGE, transferred to nitrocellulose membranes using the Trans-Blot Turbo transfer system (BIO-RAD) before blocking and incubating with 1° antibodies for the indicated protein overnight at 4 °C with shaking. The following day, membrane samples were washed, and 2° antibodies applied for 1 h at ambient temperature with shaking, washed again and imaged on a LI-COR Odyssey imager, saved as .TIFF files, and converted to presentation format with Photoshop and Illustrator software (Adobe Creative Suite).

### FLAG Immunoprecipitations

Cells were transfected with the given FLAG-tagged construct as defined in the cell culture section of methods. Collected cell pellet samples were lysed by sonication and clarified by centrifugation at 1000 x *g* for 10 min at 4 °C. Aliquots from the clarified lysate samples were combined with ANTI-FLAG M2 affinity beads (Millipore SiGMa) in Pierce Micro-Spin columns (ThermoFisher Scientific) and the remaining lysate was used to prepare input samples for Western analysis. The Micro-Spin samples were incubated at 4 °C overnight, eluted, washed twice with one column volume of PBS, and proteins eluted with 3x-FLAG peptide followed by Western blot analysis.

### LDH cell death assay

Supernatant samples were collected directly from experiments and analyzed by CyQUANT LDH cytotoxicity assay (ThermoFisher Scientific) following the manufacturers recommendations in black 384-well microplates. The samples were analyzed on a Cytation 5 multimodal plate reader (BioTek) and exported to spreadsheet software for statistical analysis. The data was visualized using GraphPad PRISM 9.2.0 Unless otherwise indicated, data represent the mean value ± SEM for 3 biological replicates.

### IL-1β/IL-18 ELISA assays

Supernatant samples were collected and pre-diluted 5-fold before quantification of cytokines by ELISA assays targeting IL-1β (R&D systems) or IL-18 (Abcam) following the manufacturers recommendations. The samples were analyzed on a Cytation 5 multimodal plate reader (BioTek) or a Tecan Infinite M1000 Pro plate reader and the data exported to spreadsheet software for statistical analysis. Data was visualized using GraphPad PRISM 9.2.0 Unless otherwise indicated, data represent the mean value ± SEM for at least 3 biological replicates.

### Microscopy

Cells were plated as defined in the cell culture section of methods and experiments performed as described within the text. At the conclusion of the experiment, cell nuclei were stained with Hoechst 33342 dye (0.8 µM) and imaged on a Zeiss Axio Observer.Z1 inverted wide-field microscope using 40×/0.95NA air objective. For each replicate well, 8-12 field positions were imaged for bright-field, DAPI, TRITC, and FITC channels and the raw data saved to .czi format for downstream analysis in ImageJ/FIJI using custom macros to calculate the total number of GFP-ASC specks and individual cell nuclei. ASC specks quantitation was calculated as the ratio of the number of GFP-ASC foci over the total number of nuclei within the field and expressed as a percentage. Data presented represent the mean value from the average of all field positions within the biological replicate. Representative images from each condition were produced within ImageJ/FIJI.

### Mitochondrial ROS analysis

3×10^4^ *CASP1*^-/-^ N/TERT-1 cells were seeded per well of a black, clear bottom 96-well microplate and cultured at 37 °C overnight. The following day, cellular ROS was evaluated for oxidants/antioxidants using the DCFDA cellular ROS assay (Abcam) following manufacturers recommendations. Briefly, culture media was aspirated, and the cells washed with 1X dilution buffer (DB) before 45 min incubation with the DCFDA reagent at 37 °C protected from light. Cells were further washed with DB, aspirated again, followed by the introduction of the indicated compounds as a 150 µL addition in PBS. The microplate was then immediately placed within a Cytation 5 multi-modal plate reader (BioTek) and the fluorescence (ex/em: 485/535 nm) read over a kinetic time course. The data was exported to spreadsheet software for statistical analysis and visualized using GraphPad PRISM 9.1.2. Unless otherwise indicated, data represent the mean ± SEM for 6 replicates.

### Cell viability measurements

2×10^3^ per well were plated in white 384-well clear-bottom tissue culture microplates (Corning) using an EL406 Microplate Washer/Dispenser (BioTek) to 25 µL final volume with the appropriate cell culture medium. The indicated compounds were introduced analytically using a CyBio pintool and the plates left to incubate at 37 °C prior to the addition of the CellTiter-Glo reagent (Promega, G7573) according to the manufacturer’s recommendations. Assay plates were placed on an orbital sharker for 2 min and incubated at 25 °C for 10 min before luminescence was read on a Cytation 5 multimodal plate reader (BioTek). The data was exported to spreadsheet software for analyses and visualized with GraphPad PRISM software (v 9.2.0).

## References and Notes

1. A. D’Osualdo et al., CARD8 and NLRP1 undergo autoproteolytic processing through a ZU5-like domain. PLoS One 6, e27396 (2011).

2. J. N. Finger et al., Autolytic proteolysis within the function to find domain (FIIND) is required for NLRP1 inflammasome activity. J. Biol. Chem. 287, 25030–25037 (2012).

3. B. C. Frew, V. R. Joag, J. Mogridge, Proteolytic processing of Nlrp1b is required for inflammasome activity. PLoS Pathog. 8, e1002659 (2012).

4. A. Sandstrom et al., Functional degradation: A mechanism of NLRP1 inflammasome activation by diverse pathogen enzymes. Science, (2019).

5. A. J. Chui et al., N-terminal degradation activates the NLRP1B inflammasome. Science 364, 82–85 (2019).

6. L. R. Hollingsworth et al., DPP9 sequesters the C terminus of NLRP1 to repress inflammasome activation. Nature 592, 778–783 (2021).

7. M. Huang et al., Structural and biochemical mechanisms of NLRP1 inhibition by DPP9. Nature 592, 773–777 (2021).

8. K. S. Robinson et al., Enteroviral 3C protease activates the human NLRP1 inflammasome in airway epithelia. Science 370, (2020).

9. B. V. Tsu et al., Diverse viral proteases activate the NLRP1 inflammasome. bioRxiv, 2020.2010.2016.343426 (2020).

10. S. Bauernfried, M. J. Scherr, A. Pichlmair, K. E. Duderstadt, V. Hornung, Human NLRP1 is a sensor for double-stranded RNA. Science, (2020).

11. M. C. Okondo et al., DPP8 and DPP9 inhibition induces pro-caspase-1-dependent monocyte and macrophage pyroptosis. Nat. Chem. Biol. 13, 46–53 (2017).

12. M. C. Okondo et al., Inhibition of Dpp8/9 Activates the Nlrp1b Inflammasome. Cell Chem Biol 25, 262–267 e265 (2018).

13. D. C. Johnson et al., DPP8/DPP9 inhibitor-induced pyroptosis for treatment of acute myeloid leukemia. Nat. Med. 24, 1151–1156 (2018).

14. F. L. Zhong et al., Human DPP9 represses NLRP1 inflammasome and protects against autoinflammatory diseases via both peptidase activity and FIIND domain binding. J. Biol. Chem. 293, 18864–18878 (2018).

15. K. Gai et al., DPP8/9 inhibitors are universal activators of functional NLRP1 alleles. Cell Death Dis. 10, 587 (2019).

16. A. R. Griswold et al., DPP9’s Enzymatic Activity and Not Its Binding to CARD8 Inhibits Inflammasome Activation. ACS Chem. Biol. 14, 2424–2429 (2019).

17. A. J. Chui et al., Activation of the CARD8 Inflammasome Requires a Disordered Region. Cell Rep 33, 108264 (2020).

18. H. Sharif et al., Dipeptidyl peptidase 9 sets a threshold for CARD8 inflammasome formation by sequestering its active C-terminal fragment. Immunity, (2021).

19. D. A. Bachovchin, NLRP1: A jack of all trades, or a master of one? Mol. Cell in press, (2021).

20. Y. He, M. Y. Zeng, D. Yang, B. Motro, G. Nunez, NEK7 is an essential mediator of NLRP3 activation downstream of potassium efflux. Nature 530, 354–357 (2016).

21. H. Shi et al., NLRP3 activation and mitosis are mutually exclusive events coordinated by NEK7, a new inflammasome component. Nat. Immunol. 17, 250–258 (2016).

22. H. Sharif et al., Structural mechanism for NEK7-licensed activation of NLRP3 inflammasome. Nature 570, 338–343 (2019).

23. D. P. Ball et al., Caspase-1 interdomain linker cleavage is required for pyroptosis. Life Sci Alliance 3, (2020).

24. T. Wang et al., Identification and characterization of essential genes in the human genome. Science 350, 1096–1101 (2015).

25. J. Snider, G. Thibault, W. A. Houry, The AAA+ superfamily of functionally diverse proteins. Genome Biol. 9, 216 (2008).

26. J. A. Duncan et al., Cryopyrin/NALP3 binds ATP/dATP, is an ATPase, and requires ATP binding to mediate inflammatory signaling. Proc. Natl. Acad. Sci. U. S. A. 104, 8041–8046 (2007).

27. R. C. Coll et al., MCC950 directly targets the NLRP3 ATP-hydrolysis motif for inflammasome inhibition. Nat. Chem. Biol. 15, 556–559 (2019).

28. Z. Hu et al., Crystal structure of NLRC4 reveals its autoinhibition mechanism. Science 341, 172–175 (2013).

29. K. C. Liao, J. Mogridge, Activation of the Nlrp1b inflammasome by reduction of cytosolic ATP. Infect. Immun. 81, 570–579 (2013).

30. J. Chavarria-Smith, P. S. Mitchell, A. M. Ho, M. D. Daugherty, R. E. Vance, Functional and Evolutionary Analyses Identify Proteolysis as a General Mechanism for NLRP1 Inflammasome Activation. PLoS Pathog. 12, e1006052 (2016).

31. F. L. Zhong et al., Germline NLRP1 Mutations Cause Skin Inflammatory and Cancer Susceptibility Syndromes via Inflammasome Activation. Cell 167, 187–202 e117 (2016).

32. S. B. Drutman et al., Homozygous NLRP1 gain-of-function mutation in siblings with a syndromic form of recurrent respiratory papillomatosis. Proc. Natl. Acad. Sci. U. S. A. 116, 19055–19063 (2019).

33. S. Grandemange et al., A new autoinflammatory and autoimmune syndrome associated with NLRP1 mutations: NAIAD (NLRP1-associated autoinflammation with arthritis and dyskeratosis). Ann. Rheum. Dis. 76, 1191–1198 (2017).

34. S. L. Masters et al., NLRP1 inflammasome activation induces pyroptosis of hematopoietic progenitor cells. Immunity 37, 1009–1023 (2012).

35. C. Y. Taabazuing, A. R. Griswold, D. A. Bachovchin, The NLRP1 and CARD8 inflammasomes. Immunol. Rev., (2020).

36. W. H. Watson et al., Redox potential of human thioredoxin 1 and identification of a second dithiol/disulfide motif. J. Biol. Chem. 278, 33408–33415 (2003).

37. M. Wang, C. J. Herrmann, M. Simonovic, D. Szklarczyk, C. von Mering, Version 4.0 of PaxDb: Protein abundance data, integrated across model organisms, tissues, and cell-lines. Proteomics 15, 3163–3168 (2015).

38. Y. Fan, M. Makar, M. X. Wang, H. W. Ai, Monitoring thioredoxin redox with a genetically encoded red fluorescent biosensor. Nat. Chem. Biol. 13, 1045–1052 (2017).

39. J. M. Herrmann, U. Jakob, Special issue: redox regulation of protein folding. Preface. Biochim. Biophys. Acta 1783, 519 (2008).

## Supplementary references

40. M. A. Dickson et al., Human keratinocytes that express hTERT and also bypass a p16(INK4a)-enforced mechanism that limits life span become immortal yet retain normal growth and differentiation characteristics. Mol. Cell. Biol. 20, 1436–1447 (2000).

41. Y. Perez-Riverol et al., The PRIDE database and related tools and resources in 2019: improving support for quantification data. Nucleic Acids Res. 47, D442–D450 (2019).

42. T. Wang et al., Identification and characterization of essential genes in the human genome. Science 350, 1096–1101 (2015).

43. J. Jumper et al., Highly accurate protein structure prediction with AlphaFold. Nature 596, 583–589 (2021).

